# Mechanistic Insights into Adaptive Binding of Vitamins and Hormones at Hotspot-Enriched Interfaces of the Synaptotagmin 1 C2AB Domain

**DOI:** 10.64898/2026.10.18.760018

**Authors:** Suvojit Hazra, Basant K. Tiwary, Alok Ghosh Chaudhuri, Nilkanta Chakrabarti

**Affiliations:** CPEPA-UGC Centre for “Electro-physiological and Neuro-imaging studies including Mathematical Modelling”, University of Calcutta, Kolkata, West Bengal, India; Department of Physiology, University of Calcutta, Kolkata, West Bengal, India; Department of Bioinformatics, School of Life Sciences, Pondicherry University, Pondicherry, India; Department of Physiology, Vidyasagar College, Kolkata, West Bengal, India

**Keywords:** Synaptotagmin 1 C2AB domain, vitamins and hormones, *In silico* binding and ranking

## Abstract

The present *in silico* study provided the adaptive binding of seven physiologically active molecules, including four non-peptide hormones (cortisol, estradiol, triiodothyronine, melatonin) and three vitamins (methyltetrahydrofolate, pyridoxal phosphate, thiamine), alongside argireline as a benchmark, at hotspot-enriched interfaces of full C2AB-domain of Synaptotagmin 1 (Syt1), using molecular docking study and molecular dynamics (MD) simulations. Linear correlations of MD-features and nonlinear KPCA-IG–based ranking of features and ligands yielded deeper mechanistic insights. All ligands exhibited spontaneous binding with strong negative ΔG_binding_ values. Fifteen hotspots, clustered across the C2A, C2B, and C2AB–interlink regions, were dominated by hydrophobic and van der Waals interactions. Several hotspots and non-hotspots corresponded to Syt1 residues that are reported to link with Ca^2+^-dependent phenomena and membrane vesicular fusion. Solvent-excluded volume (V_SE_) analyses showed that local hotspot burial and global desolvation together influenced protein adaptability, distinguishing ligands with similar hotspots but divergent thermodynamics. Linear correlations established ΔG_binding_ as an indicator of hydrophobic hotspot burial. KPCA-IG identified ΔV_SE_ and RMSD as dominant mechanistic features capturing cooperative induced-fit responses. Pyridoxal phosphate, melatonin and triiodothyronine emerged as mechanistically top adaptive binders, involving adjustable desolvation, hotspot expansion, optimal flexibility and pose retention. Methyltetrahydrofolate and cortisol showed intermediate burial-dominated cooperativity. Argireline, thiamine and estradiol exhibited ligand-specific burial-dominated, pose-locked and adaptability-limited bindings. Overall, our study demonstrates that desolvation control, induced-fit adaptability and structural flexibility, not ΔG_binding_ alone, govern Syt1-ligand binding, revealing Syt1 as a versatile presynaptic target for modulation by these bioactive molecules.

**Graphical Abstract:** 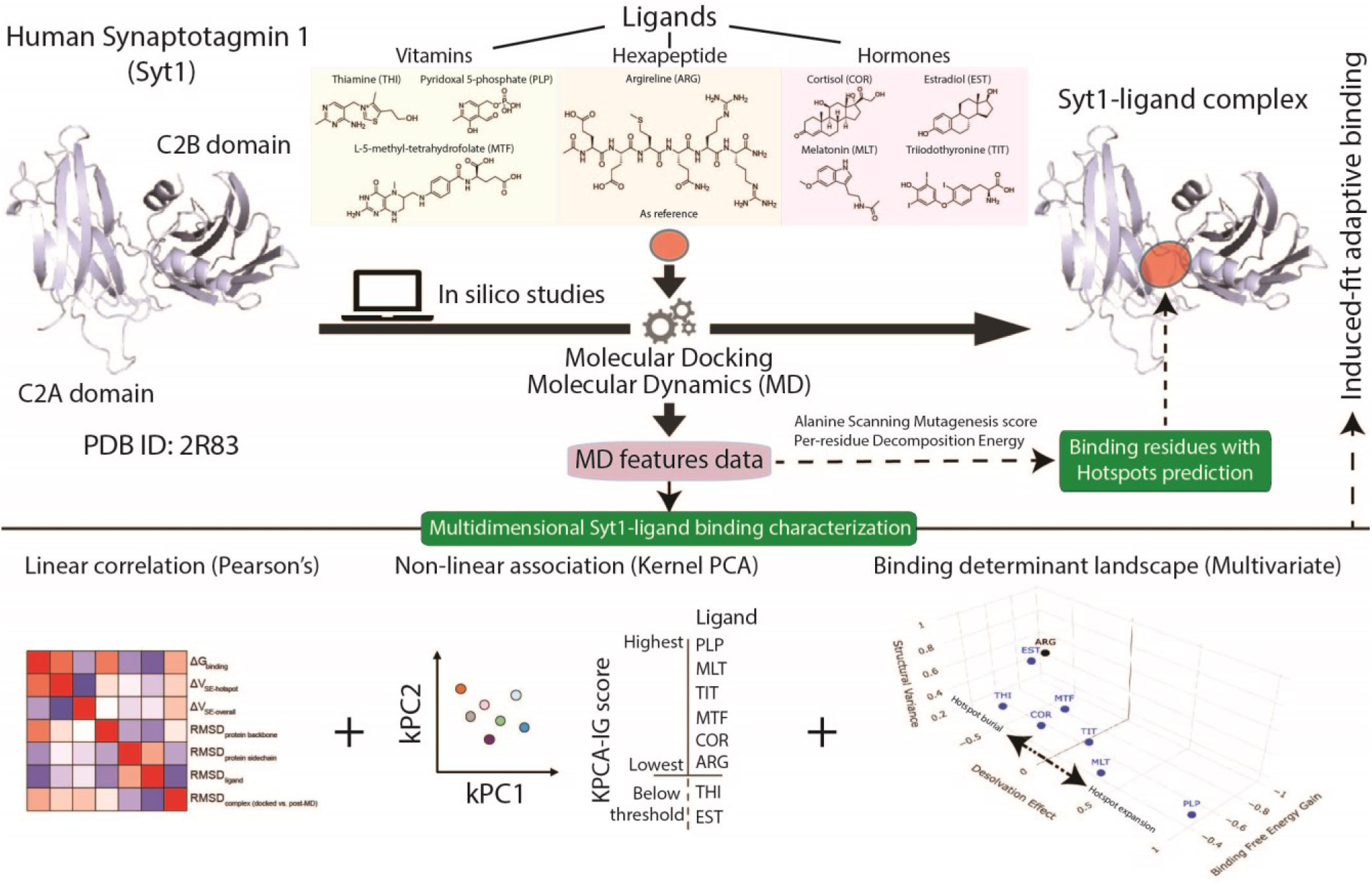

**Insight Box:** *In silico* studies using molecular docking and molecular dynamics simulations show, for the first time, that the Synaptotagmin-1 (Syt1) C2AB domain binds spontaneously and favourably to vitamins (B1, B6, and active-form B9) and hormones (cortisol, estradiol, triiodothyronine, and melatonin) at hotspot-enriched C2A, C2B, and inter-domain interfaces. Hotspot energetic decomposition, solvent-excluded volume, and RMSD analyses indicate that ligand recognition is governed by cooperative hydrophobic van der Waals packing, desolvation-driven solvent reorganization, and conformational adaptability. Kernel-based nonlinear multivariate ranking reveals that induced-fit flexibility and solvent reorganization are key determinants of productive binding, distinguishing adaptive, passive, and pose-locked binding modes. This thermodynamic-structural-solvation framework indicates that bioactive ligand binding to Syt1 may modulate calcium-dependent vesicle fusion and guide presynaptic drug discovery.

## 1. Introduction

‘Synaptotagmin 1’ (Syt1), a Ca^2+^-sensor protein of presynaptic vesicular membrane, regulates fast synchronous neurotransmitter release ^1,2^. Human Syt1 (422 residues) comprises an intravesicular N-terminal region (1–57), a transmembrane helix (58–80), and a cytoplasmic C-terminal domain (81–422) containing two homologous C2 domains namely C2A (142–263) and C2B (273–422), sharing similarity in both primary sequences (∼31% identical and ∼56% similar) and three-dimensional (3D) structure ^3^. Syt1 has two linker regions, one (residues 81-141) between the transmembrane and cytosolic domains and another flexible linker (residues 264-272) between the C2A and C2B domains ^4,5^. Syt1 binds to five Ca^2+^ ions, three in C2A and two in C2B, via the acidic side chains of conserved aspartate residues located on loop 1 and loop 3 of each C2 domain ^1,2^. The C2B domain also contains two other sites for binding SNARE (soluble N-ethylmaleimide-sensitive fusion protein attachment receptor) complex (SNARE-complex) via arginine residues ^6^ and phosphatidylinositol-4,5-bisphosphate (PIP2) of pre-synaptic plasma membrane via lysine residues ^5,7^.

Syt1 regulates vesicle fusion to the plasma membrane through coordinated, yet distinct actions of its C2A and C2B domains ^5^. It begins with anchoring and priming through (a) Syt1–oligomerization via lysine-rich juxtamembrane linker in vesicle ^4,8^, (b) a tripartite, Syt1(C2B)–(SNARE-complex)**–**complexin assembly formation ^1,6^, (c) active zone organization by chaperon-mediated multidomain protein complex and vesicular tethering via its Rab proteins to Ca²⁺ channels on plasma membrane ^2,9^, (d) C2B-mediated dual bridges formation with SNARE–complexin and syntaxin1–SNAP25 by two separate C2B, while their polybasic regions anchor to PIP2 on the inner leaflet of the plasma membrane ^5,7^, (e) C2B binding to both membrane lipid and SNARE-complex allowed by flexible C2A–C2B linker ^4,8^. Next, action potential–evoked Ca²⁺ influx triggers sequential events including (a) C2 domains reorientation inserting C2A and C2B loops into vesicular membrane and plasma membrane via phosphatidylserine (PS) and PIP2 respectively, with C2B remaining still attached to SNARE-complex ^6,7^, (b) tension generation in the flexible (C2A-C2B)-interlink, causing SNARE-complex zippering and extension of juxtamembrane linkers of synaptobrevin and syntaxin, thereby reducing membrane distance and lowering fusion energy barrier ^2,6^. Such molecular organization involves a phase transition of condensates from a liquid state during anchoring and priming processes to a gel state that stabilizes the fusion machinery, which is reversed by Ca²⁺ influx to enable vesicle fusion ^1,4,8^. Importantly, fusion pore stability depends on electrostatic C2B–PIP2 interactions that cluster local lipids and stabilize the inherently energetically unfavourable expanded fusion pore, without which pores collapse and exocytosis fails ^10^.

Syt1 has been identified as a potential drug target to control neurotransmitter release. Sevoflurane (surgical anaesthetic) can bind to Syt1–C2A domain and modulates cognition ^11^. 6-mer palmitoylated peptide (analgesic) and its N-acetylated derivative (namely argireline, an anti-wrinkle agent) can block Syt1(C2B)-SNARE complex formation and inhibit the α-calcitonin gene-related peptide (α-CGRP) exocytotic release ^12^. *In silico* study indicates that argireline can also bind to the interfacial cleft of Syt1–C2AB structure ^13^. Studies link genetic variants of Syt1 to neurodevelopmental disorders ^14^. Folate delivery to brain by high-affinity membrane transporters (FRα, PCFT) potentiates one-carbon metabolism and neurotransmitter synthesis associated with cognition and mood ^15,16^. Multiple studies indicate that neurotropic/neuromodulatory bioactive molecules including hormones like triiodothyronine ^17^, estradiol ^18^, cortisol ^19^ and melatonin ^20^ and, vitamins viz. L-5-methyl-tetrahydrofolate ^21^, pyridoxal 5-phosphate ^22^ and thiamine ^23^ are either localized at presynaptic terminals or involved in neurotransmission, collectively modulating cognitive functions. Based on these observations, we explored the potential interactions of these bioactive molecules and Syt1 through molecular docking and simulation studies. In this study, we used the native Syt1–C2AB structure instead of its isolated fragments to better explore its complete functional attributes. For comparison, argireline binding to Syt1–C2AB had been considered as a positive control.

## 2. Materials and methods

### 2.1. Identification and preparation of structure of human Syt1

The available 3D crystal structure of the cytosolic C2AB domain of human Syt1 (PDB ID 2R83; resolution 2.7Å) ^3^ was retrieved from protein data bank (RCSB-PDB) in .pdb file format. The macromolecule was exported to Autodock tools (ADT) 1.5.6 ^24^, and a grid box (dimension: 126 x 126 x 126 Å, spacing: 0.375 Å) was generated to define the molecular docking space using the graphical user interface of ADT. Structure preparation involved (i) removal of the co-crystallized ions and water molecules, (ii) adding polar hydrogens to the amino acid residues followed by the assignments of united atom Gasteiger charges, solvation parameter and fragmental volume, and (iii) saving of the prepared files in *.pdbqt* (*.pdb* plus *q* [Gasteiger charges] and *t* [AutoDock type]) format, using ADT.

### 2.2. Ligand structure identification and preparation

The 3D structure of the eight ligand molecules were retrieved from the PubChem database ^25^. Five physicochemical properties (molecular weight [in Dalton], number of hydrogen bond donor (HBD) and acceptor (HBA), logP (octanol-water partition coefficient) and total polar surface area [tPSA; in Å^2^]) were extracted using SwissADME web-server ^26^. Using ADT (Autodock Tools), the Gasteiger partial charges and torsions were allocated, Kollman charges and polar hydrogen atoms were applied, and *.pdbqt* files of ligands were generated before molecular docking study.

### 2.3. Molecular Docking study

Autodock Vina (v1.2.1) software ^27^ was implemented for the calculation of the binding affinity of the ligands on Syt1–C2AB using processed *.pdbqt* files of protein and ligands with the global searching exhaustiveness parameter with default value 8.0 to get the best-docked conformations ^28^. The top protein-ligand complex conformation in each ligand was selected based on the lowest Autodock docking energy scores (ADS).

### 2.4. Molecular dynamics (MD) simulation study

All docked protein–ligand complexes were subjected to MD simulations using the AMBER(v18) package ^29^. Systems were employed with GAFF (General Amber Force Field) and ff99SB force fields, solvated in a TIP3P water model filled in an octahedral box (edge-length: 20 Å) and neutralized with appropriate Na⁺/Cl⁻ counterions to maintain physiological ionic strength. Energy minimization was followed by gradual heating from 0 to 310 K (37 °C) under a 2 kcal.mol^-1^.Å^-2^ restraint for 0.05 nanosecond, relaxing only water molecules and counterions. Density equilibration was conducted under the isobaric-isothermal (1 atm: 1.013 bar; 310 K) NPT ensemble in three successive phases of 0.05 nanosecond at 2 kcal.mol^-1^.Å^-2^ restraint, 0.05 ns at 1 kcal.mol^-1^.Å^-2^ restraint and 0.1 nanosecond unrestrained. A 10-nanosecond production run was performed under the NPT ensemble at 310 K (maintained by Langevin piston thermostat scheme) temperature and 1 atm (maintained by Nosé-Hoover-Langevin piston scheme) pressure using a 2 femtoseconds timestep. Long-range electrostatic interactions were treated by the Particle–Mesh Ewald method with a 12Å cut-off range. All bonds involving hydrogen atoms were constrained using the SHAKE algorithm.

Post-MD binding energy calculations were performed using the MM-GBSA (Molecular Mechanics-Generalized Born Surface Area) approach implemented in the AMBER *mmpbsa.py* module [29]. Binding free energy (ΔG_binding_), per-residue decomposition energy (PRDE) and *in silico* alanine scanning mutagenesis (ASM) score (ASM-ΔΔG) were computed from time-resolved trajectory frames. The mean energy for each nanosecond was calculated from 500 evenly spaced frames. Each frame (2 picoseconds interval) corresponded to every 1000th step of the 2 femtoseconds timestep.

Hotspot residues were identified based on ASM-ΔΔG > 2 kcal.mol^-1^ ^30,31^ and PRDE < 0 kcal.mol^-1^ corresponding to the greatest binding free energy contributions. Structural stability and conformational dynamics were assessed by calculating the best-fit Root Mean Square Deviation (RMSD) values of the ligand (RMSD_ligand_), Syt1–backbone (RMSD_protein backbone_) and Syt1–sidechain (RMSD_protein sidechain_) and as well as the distances between each ligand and the Cα atoms of hotspot residues. These quantities were evaluated for 5000 frames sampled at 2 picoseconds intervals over the 10 nanoseconds simulation using Kabsch’s algorithm implemented in the AMBER *cpptraj* module ^32^. Notably, MD simulations of the apo (ligand-free; wild) Syt1 protein were performed under identical conditions and for the same simulation length (10 ns) as those used for the protein–ligand complexes for comparison during measurement of RMSD values of protein (backbone and side chain).

Two-dimensional protein–ligand molecular interaction images were prepared using LigPlot+ ^33^. Superimposed docked and post–MD poses for each complex (RMSD_complex (docked vs. post–MD)_) were generated in PyMOL ^34^ using the super command, which applies the Kabsch least-squares fitting algorithm with an RMSD cutoff of 2.0 Å. Three-dimensional protein–ligand molecular interactions images of ligands with Syt1 hotspot residues in pre-(docked) and post–MD conformations were prepared with PyMOL.

### 2.5. Dynamic volume change analysis

Protein volume changes upon ligand binding were analyzed using ProteinVolume1.3 program ^35^ which employs a flood-fill technique to calculate molecular surface based on atomic coordinates from the protein structure (.pdb) file and then filling it with multiple 0.02 Å probes. The solvent-excluded volume (V_SE_) was computed from the sum of probe volumes using the atomic radius from all-atom Bondi radii set, while Van der Waals volume (V_vdW_) incorporated an additional check to ensure probes lie within atomic V_vdW_ radii. The void volume (V_Void_) was quantified [(V_Void_)=(V_SE_)−(V_vdW_)] as intramolecular void spaces due to imperfect amino acid packing in the protein structure.

Default settings with 0.04 Å starting probe, 0.02 Å ending probe and 0.1 Å surface probe minimum distance, were used to calculate V_SE_ of binding site pocket and total volume of the each Syt1–ligand complex, as well as the input Syt1 structure. The starting probe size was set with the initial radius, which became half on collisions with atoms or surface probes until reaching the ending probe size that specified the minimum radius for all probes, ensuring computational accuracy. The surface probe minimum distance was set with the closest spacing between surface probes, increasing of it reduced probe density, calculation, and processing time. The change in total solvent-excluded volume (ΔV_SE-overall_) upon MD simulation was measured by subtracting the V_SE-overall_ of input Syt1 structure from V_SE-overall_ of Syt1–ligand complexes. An analogous calculation was performed for individual hotspot residues to obtain ΔV_SE-hotspot_.

### 2.6. Statistical estimations

Energetic (ΔG_binding_, PRDE) and structural (RMSDs and ligand–hotspot centroid distances) features were computed as time-averaged quantities and are reported as mean ± SD, with SD capturing temporal variability in energetic heterogeneity and conformational stability, respectively, along the MD trajectory. Pose RMSD (docked vs. post-MD complexes) and V_SE_ values for the hotspot and overall protein were reported as absolute quantities, as these features represented cumulative geometric properties rather than stochastic variables for statistical averaging. The Pearson’s correlation coefficient (r) values were used to determine the strength of relationships among features, categorized as ‘very strong’ (0.8–1.0), ‘strong’ (0.6–0.79), and ‘moderate’ (0.4–0.59) correlations ^36^. The ‘Kernel principal component analysis (KPCA) with Interpretable Gradient’ (KPCA-IG) was employed to derive data-driven importance scores for identifying influential variables (features and ligands). KPCA, a non-linear extension of standard PCA, was performed using the radial basis function kernel in the R package ‘kpcaIG’. The most relevant variables were extracted based on scores computed from the partial derivatives of the kernel computation ^37^.

## 3. Results

### 3.1. Structural details of human Syt1

The crystal structure of human Synaptotagmin 1 (UniProt ID: P21579) comprised 279 residues (E140-A418), encompassing the C2A domain (L142–Q263 residues), a short linker (S264–K272 residues), and the C2B domain (L273–A418 residues).

### 3.2. Physicochemical properties of ligands

Scaled physicochemical profiles (0–1; **Fig. 1A**) showed that all eight bioactive ligands (vitamins and non-peptide hormones) satisfied key properties with threshold values of MW (150-500 Dalton), logP (−0.7 to +5.0), HBD (≤ 5), HBA (≤ 10), tPSA (20-130 Å²). Borderline deviations were observed for MTF (tPSA: 202.77 Å²) and TIT (MW: 650.97 Dalton). ARG was an exception, exhibiting markedly high MW (888.99 Dalton), HBD/HBA (14/14) and tPSA (489.48Å²), together with very low logP (-6.32), consistent with a large, highly polar ligand. Linear correlation analyses (**Fig. 1C**) showed that HBD, HBA, and tPSA values were positively correlated to each other but negatively correlated with logP, reflecting chemically consistent trends where the former three indicated hydrophilicity and the latter one lipophilicity.

**Fig. 1:**
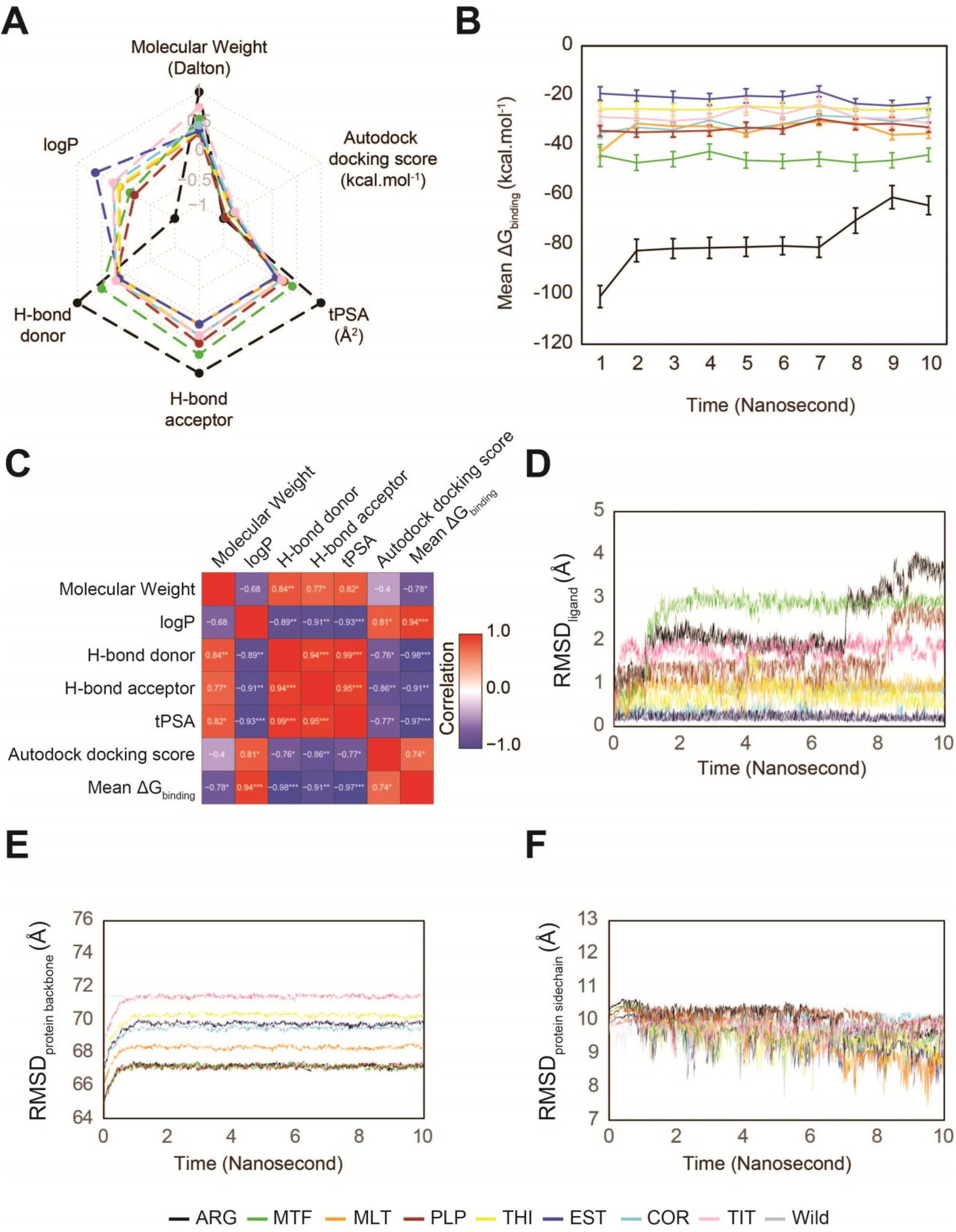
Summary of binding properties of ligands to Syt1-C2AB structure. **(A)** Radar plot shows normalized values (0 and 1) of Autodock scores and five physicochemical properties of eight ligands, providing an overview of parameters used for the selection of ligands. **(B)** MM-GBSA derived binding free energy (ΔG_binding_) values (kcal.mol^-1^) calculated over the full MD trajectories are plotted as mean±SD, representing relative binding affinities of ligands to Syt1. **(C)** Heatmap of Pearson’s correlation coefficient (*r*) values between physicochemical properties, Autodock scores and ΔG_binding_ values shown in Fig. A and B, illustrating linear correlations among parameters. The statistical significances (*** for p < 0.001, ** for p < 0.01, * for p < 0.05) of *r* values are displayed in a continuous colour gradient of blue, white and red corresponding to *r* = - 1, 0, 1, respectively. **(D-F)** Plots of time evolution of RMSD values (Å) for ligands **(D)**, protein backbone **(E)**, and protein sidechain **(F)** during MD simulations trajectories, comparing ligand-bound complexes and apo protein (wild) to assess pose stability, global fold integrity and local conformational flexibility, respectively.

### 3.3. Molecular docking study

Molecular docking identified ARG as the highest-affinity ligand (ADS: −7.6 kcal.mol^-1^), followed by MTF (−7.3), PLP (−7.2), THI (−6.4), EST (−6.1), MLT and COR (both −5.9), and TIT (−5.8). Normalised ADS values (0–1) were visualised in a radar plot together with five key physicochemical parameters (**Fig. 1A**). Linear correlation analysis (**Fig. 1C**) showed that ADS correlated positively with logP and negatively with HBD, HBA, and tPSA, indicating that higher lipophilicity favours stronger docking affinity via hydrophobic interactions.

### 3.4. Molecular dynamics study and binding mode analysis

#### 3.4.1. Binding free energy (ΔG_binding_) analysis of Syt1–Ligand Complexes

All ligand binding showed negative values of the ΔG_binding_ scores (kcal.mol^-1^) with differential binding affinities to Syt1 throughout the MD trajectories (**Fig. 1B**). Negative ΔG_binding_ values indicate thermodynamically favourable and spontaneous protein–ligand binding. Accordingly, average binding affinity ranked as ARG>MTF>MLT>PLP>COR>TIT>THI>EST (**Table 1**). Linear correlation analysis (**Fig. 1C**) showed that ΔG_binding_ correlated positively with ADS and logP and negatively with HBD, HBA, and tPSA, indicating that more lipophilic ligands with higher docking scores form thermodynamically stronger and hydrophobically driven interactions with Syt1.

**Table 1.** Binding free energies and structural stability metrics from molecular dynamics simulations. Mean binding free energy (ΔG_binding_) and RMSD values of protein backbone, protein side-chain and ligand are reported as mean ± SD, averaged over windows across 10 nanosecond trajectories for each ligand-protein complex. RMSD metrics capture global backbone stability, local sidechain adaptability, and ligand pose stability, respectively.

| Ligand | $\Delta G_{\text{binding}}$ (kcal.mol<sup>-1</sup>) | RMSD<sub>protein backbone</sub> (Å) | RMSD<sub>protein sidechain</sub> (Å) | RMSD<sub>ligand</sub> (Å) |
| --- | --- | --- | --- | --- |
| ARG | $-78.30 \pm 11.20$ | $66.34 \pm 0.28$ | $10.05 \pm 0.36$ | $2.31 \pm 0.74$ |
| MTF | $-45.37 \pm 1.48$ | $66.36 \pm 0.29$ | $9.66 \pm 0.35$ | $2.65 \pm 0.64$ |
| MLT | $-33.74 \pm 3.70$ | $67.59 \pm 0.32$ | $9.51 \pm 0.61$ | $0.91 \pm 0.15$ |
| PLP | $-32.83 \pm 1.68$ | $66.33 \pm 0.28$ | $10.13 \pm 0.23$ | $1.49 \pm 0.53$ |
| COR | $-31.19 \pm 2.45$ | $68.70 \pm 0.35$ | $9.93 \pm 0.25$ | $0.65 \pm 0.27$ |
| TIT | $-28.12 \pm 2.42$ | $70.63 \pm 0.37$ | $9.83 \pm 0.22$ | $1.78 \pm 0.22$ |
| THI | $-25.19 \pm 0.55$ | $69.54 \pm 0.39$ | $9.54 \pm 0.34$ | $0.70 \pm 0.22$ |
| EST | $-21.02 \pm 1.87$ | $69.03 \pm 0.39$ | $9.48 \pm 0.43$ | $0.21 \pm 0.06$ |

#### 3.4.2. RMSD of ligands and protein

RMSD trajectories of the ligands (**Fig. 1D**), Syt1 backbone (**Fig. 1E**), and side chains (**Fig. 1F**) remained <4 Å, indicating stable docking poses and overall complex stability for each case of study throughout the MD simulation trajectories. The analyses of average values of RMSDs revealed ligand-dependent modulation of protein stability and binding dynamics (**Table 1**). MTF exhibited the highest ligand mobility, whereas EST showed the most rigid, pose-locked binding. PLP

induced the largest relative sidechain rearrangements while maintaining low backbone RMSD, indicating pronounced local adaptation. Other ligands displayed intermediate and distinct RMSD signatures.

#### 3.4.3. Characterization of 47 binding residues

The ball-stick atomistic model of 2D interactions of post–MD complexes (**Fig. 2A-H**) depicted the putative hydrogen bonds and hydrophobic interactions between Syt1 and ligands. Cumulatively, the crystalline structure of C2AB–domain of Syt1 showed 47 residues as binding residues in interfaces of ligand bindings during MD analyses.

**Fig. 2:**
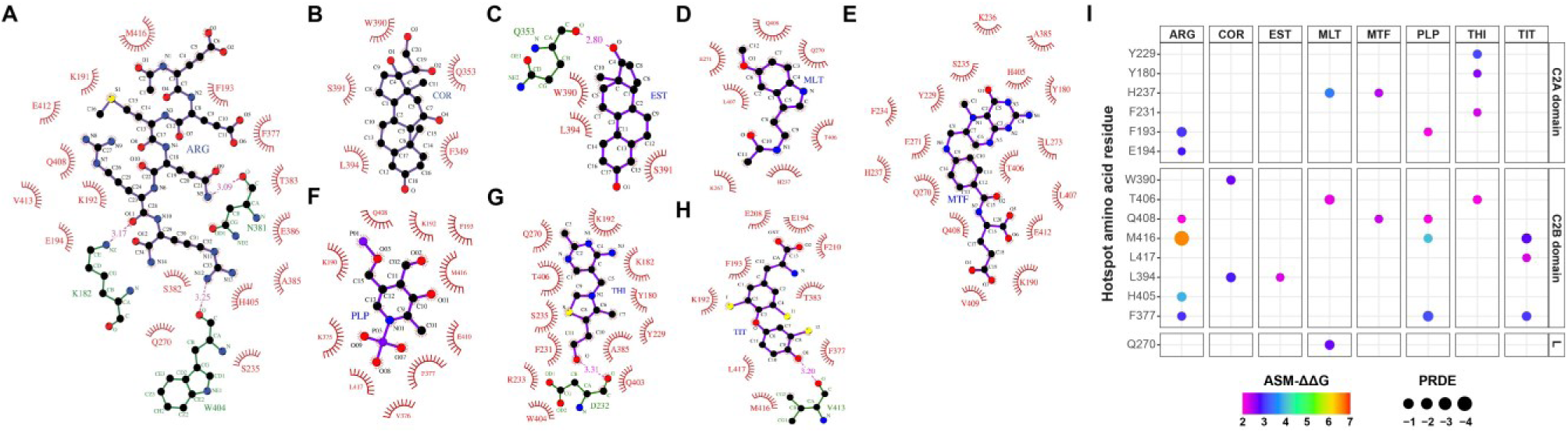
Ligand-protein interaction patterns and hotspot residues in Syt1. **(A-H)**: Atomistic 2D interaction maps of Syt1 with ligands derived from post-MD-refined complexes using LigPlot+ software, shown as ball-and-stick representations, depict persistent hydrogen bindings and hydrophobic contacts formed between Syt1 and the bound ligands ARG **(A)**, COR **(B)**, EST **(C)**, MLT **(D)**, MTF **(E)**, PLP **(F)**, THI **(G)**, and TIT **(H)**. Carbon, nitrogen, oxygen, phosphorous, sulphur/iodine atoms are shown in black, blue, red, magenta, yellow, respectively. Atomic bonds of ligand and amino acids of Syt1 involved in hydrogen bond formation are depicted in violet and green, respectively. Hydrophobic interacting amino acid residues of Syt1 are marked in red fonts and surrounding dark red radial nodes. Amino acids of Syt1 involved in hydrogen bonds are represented as green fonts. **(I):** Bubble plot summary of hotspot residues of Syt1 contributing to ligand binding, where the x-axis depicts ligands (three-letter code presented at the top of the x-axis) and the y-axis indicates hotspot residues (single-letter code and residue number presented in the left side of the y-axis), grouped by Syt1 domains (C2A, C2B) and Linker (L). Bubbles sizes adjusted in 3 to 6 pts of continuous mapping, reflect per-residue MM-GBSA decomposition energy (PRDE) scores (0 to -4 kcal.mol^-1^). Bubble colours represent ASM-ΔΔG scores (2–7 kcal.mol^-1^) using a continuous VIBGYOR color gradient.

47 residues of Syt1 appeared as binding residues in interfaces of ligand bindings. These ligand binding residues of Syt1 appeared as polar amino acids with either acidic (negatively charged viz. Aspartate: D232; Glutamate: E194, E208, E271, E386, E410, E412) or basic (positively charged viz. Lysine: K182, K190, K191, K192, K236, K267, K375; Arginine: R233; Histidine: H237, H405) or neutral (uncharged viz. Serine: S235, S382, S391; Threonine: T383, T406; Asparagine: N381; Glutamine: Q270, Q353, Q403, Q408; Tyrosine: Y180, Y229) and non-polar (*Aliphatic* viz. Alanine: A385, A386; Leucine: L273, L394, L407, L417, Methionine: M416; Valine: V376, V409, V413; and *Aromatic* viz. Phenylalanine: F193, F210, F231, F234, F349, F377; Tryptophan: W390, W404) in nature at physiological condition. Notably, histidine and tyrosine can switch between polar (positively charged) to neutral polar form and neutral polar to polar (negatively charge) form respectively.

The ligands ARG, COR, EST, MLT, MTF, PLP, THI, and TIT engaged 19(3), 5, 4(1), 7, 17, 10, 13(1), and 10(1) Syt1 residues, respectively, at their binding interfaces, with numbers in parentheses indicating hydrogen bonds and the remainder hydrophobic contacts. Total six hydrogen bond interacting residues of Syt1 appeared as W404 and K182 with ARG and THI, V413 with ARG and TIT, Q353 with COR and EST, N381 with ARG and D232 with THI. Notably, the hydrogen-bonding residues also displayed hydrophobic interactions with other ligands.

#### 3.4.4. Characterization of 15 hotspot residues

##### 3.4.4.1. Hotspot residues and their interacting ligands

15 hotspots were identified among binding residues of Syt1 with the distributions of (i) six for ARG (F193, E194, F377, H405, Q408 and M416), four for each (ii) PLP (F193, F377, Q408 and M416) and (iii) THI (Y180, Y229, F231 and T406), three for each (iv) MLT (H237, Q270 and T406) and (v) TIT (F377, M416 and L417), two for each (vi) MTF (H237 and Q408) and (vii) COR (W390 and L394), and (viii) one for EST (L394). Seven of 15 hotspots (F193, F377, H237, L394, M416, Q408, T406) recurred across ligands, indicating partial overlap of preferred binding regions. Using the standard Syt1 (human) boundaries ^3^ the hotspots appeared to be distributed across the C2A domain (Y180, F193, E194, Y229, F231, H237), linker/N-C2B edge (Q270), and C2B domain (F377, W390, L394, H405, T406, Q408, M416, L417) of Syt1 (**Fig. 2I**).

##### 3.4.4.2. ASM score of hotspot residues

All ASM-ΔΔG values (kcal.mol^-1^) for key hotspots after 10 ns of MD simulations exhibited an increase in binding free energy of > 2.0 kcal.mol^-1^, with highest value observed for M416A (6.62 kcal.mol^-1^) in the Syt1–ARG complex and lowest for F193A (2.01 kcal.mol^-1^) in the Syt1–PLP complex (**Fig. 2I**).

##### 3.4.4.3. Mean PRDE values of hotspot residues

All hotspots showed negative mean PRDE-total values (**Fig. 2I**), indicating net stabilizing contributions across complexes. The strongest and weakest stabilizing residues (mean ± SD, kcal.mol^-1^), respectively were (i) M416 (−4.70 ± 0.60) and E194 (−0.15 ± 0.83) in ARG complex, (ii) M416 (−3.38 ± 0.41) and F377 (−1.14 ± 0.38) in PLP complex, (iii) H237 (−1.49 ± 0.49) and Q408 (−0.86 ± 0.42) in MTF complex, (iv) T406 (−2.45 ± 0.31) and Q270 (−2.10 ± 0.51) in MLT complex, (v) M416 (−2.46 ± 0.39) and F377 (−1.78 ± 0.48) for TIT complex, (vi) L394 (−2.18 ± 0.85) and W390 (−1.54 ± 0.53) in COR complex, (vii) T406 (−1.23 ± 0.38) and Y180 (−0.26 ± 0.12) in THI complex. In EST complex, L394 (−0.76 ± 0.30) also showed residue-specific stabilization pattern. PRDE component analysis (**Fig. 3C**) showed that vdW energies (E_vdW_) provided the dominant stabilizing contributions, counterbalanced by polar solvation (E_ps_) penalties and consistently supported by non-polar solvation energies (E_nps_). PRDE analysis exhibited that among seven recurring hotspots, most of them showed strong ligand-dependent bindings, driven predominantly by vdW energy contributions (M416 in ARG and TIT, F193 in ARG, F377 in PLP and TIT, T406 in MLT and THI, L394 in COR, H237 in MLT) except weak bindings (Q408 in ARG, MTF, PLP). The remaining residues appeared to be ligand-specific, with most of them as weak (E194, F231, H405, L417, W390, Y229 and Y180) binders where vdW energetics were counterbalanced partially by polar solvation energies. However, another ligand-specific interaction (Q270 in MLT) represented strong binding.

**Fig. 3:**
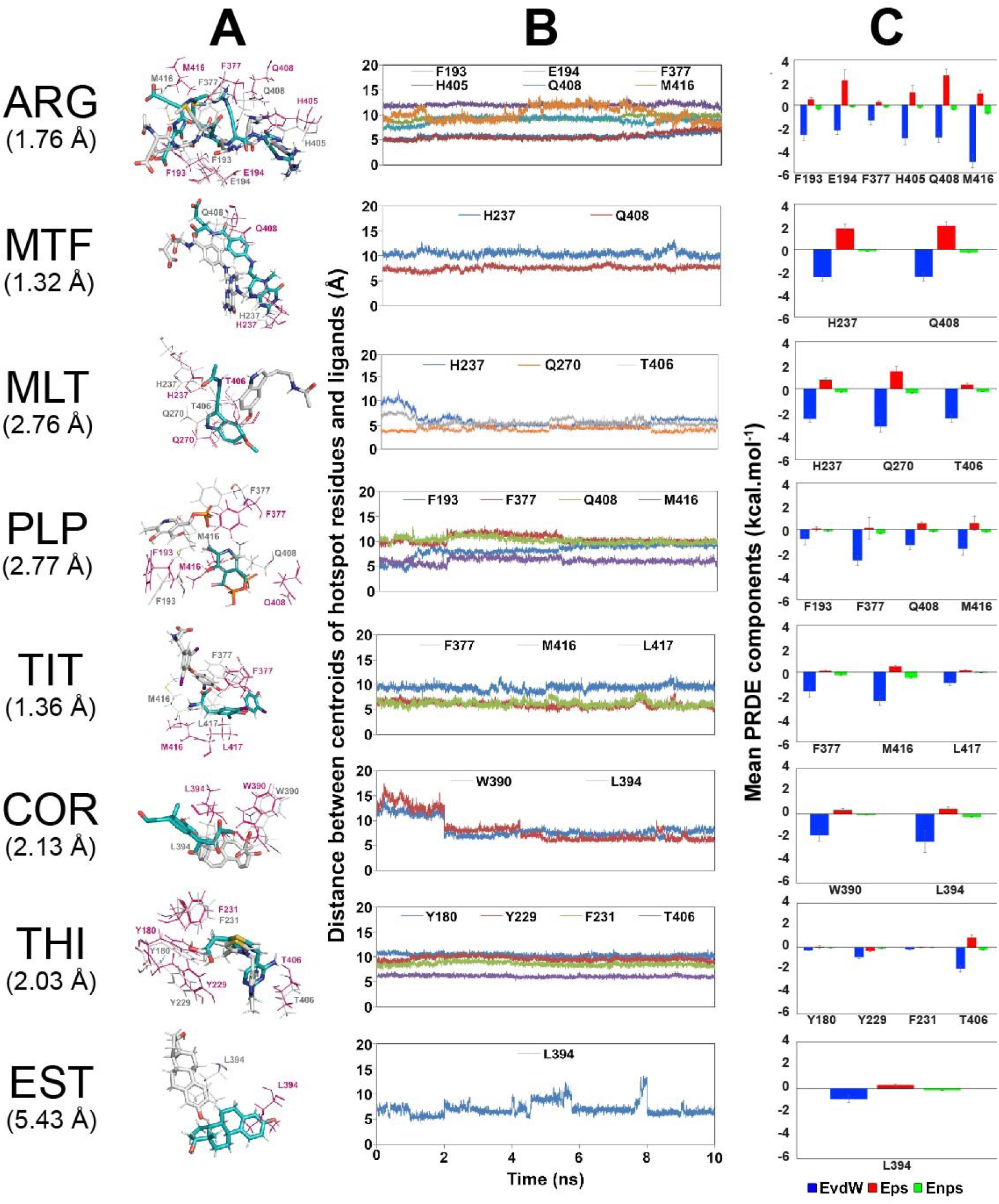
Structural stability and energetic contributions of hotspot residues during MD simulations. **(A)** Superposition of pre-MD (grey) and post-MD (cyan) binding poses of ligands and corresponding hotspot residues in Syt1-C2AB, where ligands and hotspot residues are shown as sticks and lines, respectively, illustrating pose retention and local rearrangements following dynamic equilibration. Images were generated using PyMOL with three ‘set’ commands (viz., bg_color white; set antialias,2; set ray_trace_mode,0) followed by merging the two binding poses (post-docking and post-dynamics) of eight Syt1-C2AB-ligand complexes for a better 3D representation. **(B)** Distance between Cα atom of each hotspot and the centroid of the corresponding ligand is represented in the time-scale of the MD simulation trajectories, reporting the stability of ligand-hotspot associations. **(C)** Bar plots show values (mean±SD) of van der Waals (EvdW; presented in blue colour), polar solvation (Eps; presented in red colour) and non-polar solvation (Enps; presented in green colour) energy components of hotspot residues derived from MM-GBSA decomposition at the end of MD simulations. Notably, RMSD value (Å) of each ligand with its hotspot binding poses between docked vs. post-MD structures (depicted in Fig. 3A) is indicated below the corresponding ligand labels (in the very first column).

##### 3.4.4.4. Ligand–hotspot distances and dynamic binding poses

The distances between the centroid of ligands and the Cα atom of hotspots (**Fig. 3B**) dynamically altered between <4 Å and >8 Å during MD simulation trajectories, indicating substantial conformational flexibility that led to the dynamic binding poses of both docked and post–MD ligands with hotspots (**Fig. 3A**) and stable Syt1–ligand complex formation. RMSD-complex(docked versus post–MD)-pose values (RMSD_complex (docked vs. post–MD)_) showed the lowest pose deviation for MTF (1.32 Å) and highest for EST (5.43 Å) during complex formation (**Fig. 3**).

### 3.5. Dynamic volume change analysis

In our study, five ligands (ARG, COR, EST, MTF, and THI) showed a decrease in V_SE-hotspot_ (negative ΔV_SE-hotspot_) with maximal changes (-30.644 Å^3^) in ARG and minimal change (-0.126 Å^3^) in EST. Other three ligands (TIT, MLT, PLP) showed increases in V_SE-hotspot_ (positive ΔV_SE-hotspot_) with maximal change (15.945 Å^3^) in PLP and minimal change (6.599 Å^3^) in TIT (**Fig. 4**). The Syt1–ligand complexes exhibited an increase in ΔVSE_-overall_ (positive ΔV_SE-overall_) with maximum change (384.574 Å^3^) in ARG-complex and minimum change (150.918 Å^3^) in PLP-complex (**Fig. 4**).

**Fig. 4:**
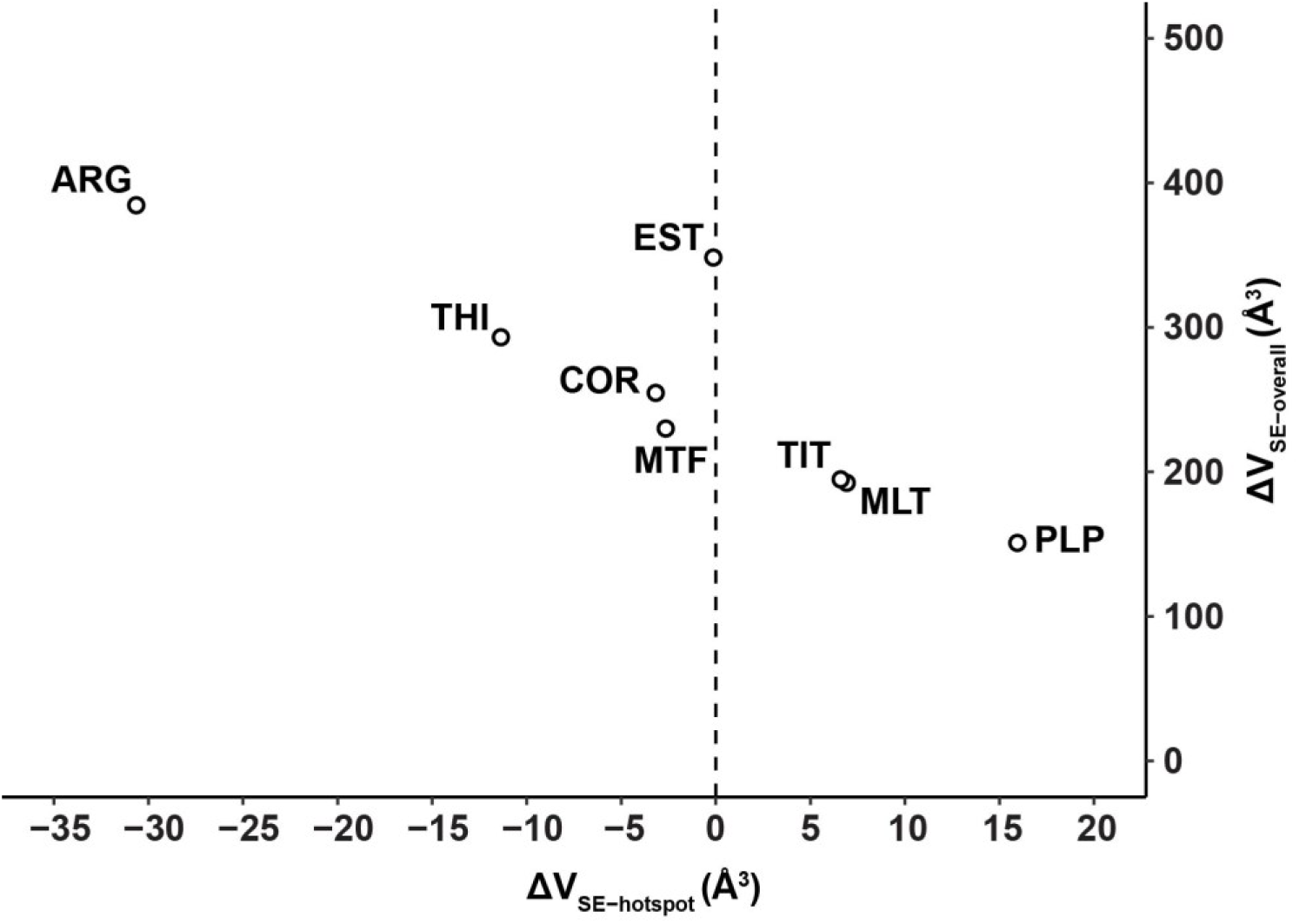
Ligand-dependent ‘solvent-excluded volume’ changes (ΔV_SE_) during Syt1 binding. Scatter plot shows ΔV_SE_ values (Å³) at the binding hotspot site (ΔV_SE_-_hotspot_) versus the overall protein (ΔV_SE_-overall) for individual ligand-Syt1 complexes. ΔV_SE_ values are calculated as the difference between the beginning and end of the MD simulation trajectories, used as geometric descriptors of ligand-induced solvent reorganizations, reporting local binding-site burial (ΔV_SE_-_hotspot_) and global protein desolvation (ΔV_SE_-_overall_) associated with complex formation.

### 3.6. Correlation analyses of features and ligands

Both linear and nonlinear correlation analyses were performed using pose-deviation values (**Fig. 3**), ΔV_SE_ values (**Fig. 4**), and the averaged values of ΔG_binding_ and RMSDs (**Table 1**). The ΔG_binding_ values (increasingly negative) showed significant (p<0.05) strong linear negative correlations (r≥0.6) with V_SE-hotspot_. Significant (p<0.05) strong linear correlations with negative magnitude were also observed for V_SE_ features comparing hotspot and whole-protein (overall) regions. Strong linear correlations (r≥0.6) exhibiting only trend-level significance (0.05<p<0.10) were identified between (a) ΔG_binding_ and RMSD_protein backbone_ as well as RMSD_ligand_, and (b) RMSD_ligand_ and RMSD_complex (docked vs. post–MD)_. Other strong linear correlations (r≥0.6) were not statistically significant (p>0.05).

KPCA score–loading biplots (**Fig. 5B**) were constructed to visualize nonlinear contributions of MD-derived features and ligands during protein–ligand binding and to provide a geometric basis for interpreting KPCA-IG score rankings by mapping variable contributions onto the kernel-transformed variance structure.

**Fig. 5:**
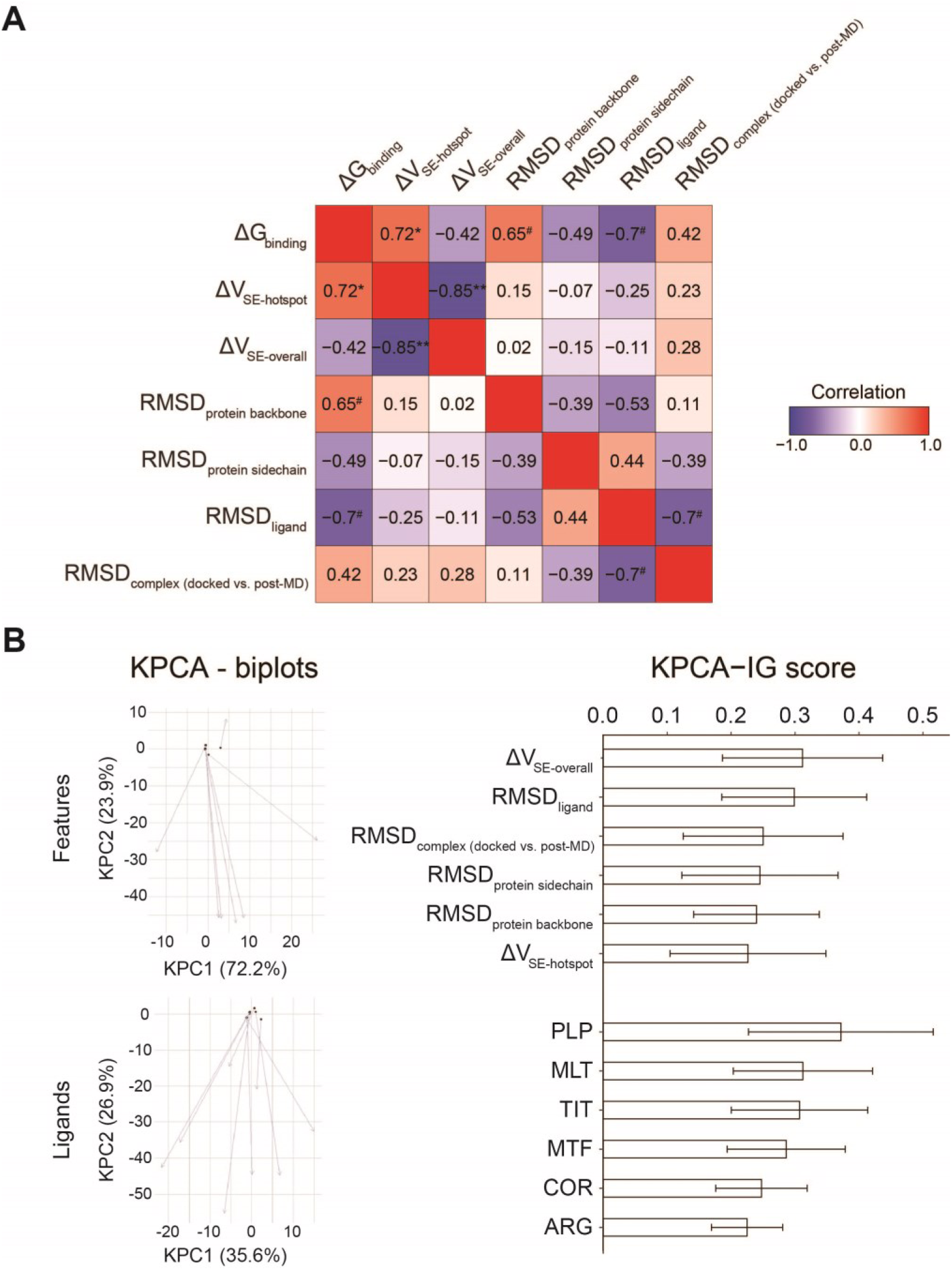
Linear correlations of MD-derived features and KPCA-IG–based nonlinear ranking of both features and ligands to Syt1 binding. **(A)** Heatmap of Pearson’s correlation coefficient (*r*) values between seven MD-derived features reports pairwise linear relationships. Numerical r values are shown with significance levels (# for p<0.10, * for p<0.05 and ** for p<0.01). Other values without having any symbol are not significant. **(B)** Kernel principal component analysis (KPCA) score-loading biplots (left panel) are constructed from the same kernel-transformed data matrix values of seven MD-derived features across eight ligand-Syt1 complexes. Nonlinear projections onto the first two kernel principal components (kPC1 and kPC2) are shown in feature space (top of left panel) and ligand space (bottom of left panel). Feature-space KPCA-biplot exhibits a strongly skewed variance distribution (kPC1: 72.2%, kPC2: 23.9%), indicating a hierarchical nonlinear organization of binding determinants dominated by a single principal axis. In contrast, ligand-space KPCA-biplot shows a more balanced variance distribution (kPC1: 35.6%, kPC2: 26.9%), reflecting multiaxial and cooperative ligand-specific binding responses. Kernel PCA-interpretable gradient (KPCA-IG) values (right panel) show the top six ranked MD-derived features and ligands, generated using the ‘kpcaIG’ R package by combining nonlinear kernel eigen decomposition with partial-derivative based attribution to quantify contributions to the dominant nonlinear variance manifold.

Feature-space KPCA revealed a strongly skewed variance distribution (kPC1: 72.2%, kPC2: 23.9%), with KPCA-IG governed by a single principal nonlinear axis defining descriptor ranking. Accordingly, KPCA-IG ranked ΔV_SE-overall_>RMSD_ligand_>RMSD_complex (docked vs. post–MD)_>RMSD_protein sidechain_>RMSD_protein backbone_>ΔV_SE-hotspot_ as the most influential descriptors, while ΔG_binding_ fell outside this dominant group.

In contrast, ligand-space KPCA showed a more balanced variance distribution (kPC1: 35.6%, kPC2: 26.9%), with KPCA-IG integrating multiaxial contributions to rank PLP>MLT>TIT>MTF>COR>ARG. Ligands outside the top ranks (THI and EST) clustered near the origin of the kPC1–kPC2 manifold and contributed minimally to the dominant nonlinear variance.

## 4. Discussion

The present *in silico* study demonstrated for the first time, that the full native Syt1–C2AB domain interacted with seven physiologically relevant neuromodulatory molecules including non-peptide hormones (cortisol, triiodothyronine, estradiol, melatonin) and vitamins (B1, B6, B9-active form). Argireline, a known ligand reported to stabilize the closed conformation of Syt1 ^13^, served as a benchmark validating computational and MD protocols. These integrative approaches enhanced system stability and binding fidelity, establishing the utility of argireline for probing Syt1 interactions with other bioactive ligands. Thermodynamically, all ligands bound spontaneously and favourably (**Fig. 1B**), with strong affinities (negative ΔG_binding_) influenced by lipophilicity (**Fig. 1A, 1C**). Binding residues (**Fig. 2A–H**) were stabilized predominantly by hydrophobic and vdW interactions across 15 hotspots spanning the C2A, C2B, and C2AB–interlink regions of Syt1, supported by ASM-ΔΔG scores (**Fig. 2I**), PRDE (**Fig. 2I, 3C**) and interfacial centroid distances between ligands and hotspot Cα atoms (**Fig. 3A-B**). RMSD (**Fig. 1D–F, 3**) and V_SE_ (**Fig. 4**) measurements provided complementary dynamic and spatial insights, respectively, at both global and binding-interface levels. Linear (**Fig. 5A**) and nonlinear (**Fig. 5B**) dynamic correlations of MD-features established the thermodynamic and mechanistic basis of Syt1–ligand interactions.

### 4.1. Ligand properties and binding forces

Small bioactive molecules (vitamins and non-peptide hormones) with favourable physicochemical properties (**Fig. 1A**) exhibited strong peptide binding potentials. ARG was represented as a non-traditional peptide binder driven by structural complementarity and binding strength. It is reported that ligand efficiency for small molecules reflects balanced enthalpy, entropy, and geometry ^38^, with stereochemical and aromatic optimization preserving pharmacokinetic integrity ^39^, while short peptides achieve interface-adaptive binding with limited entropic cost ^40^. In the present study, all ligands were subjected to molecular docking and dynamics simulations to evaluate their interactions with Syt1, considering enthalpic, entropic, and structural factors of ligand efficiency.

Our study showed that negative ADS (**Fig. 1A**) and ΔG_binding_ scores (**Fig. 1B**) were positively correlated (**Fig. 1C**), supporting efficient ligand screening and binding affinities ranking. AutoDock Vina ranks ligand poses by increasingly negative scores, but approximates thermodynamics only partially due to simplified entropy and missing electrostatic and solvation terms, with rankings sensitive to docking parameters ^28,41^. Binding free energy (ΔG_binding_) quantifies protein–ligand affinity, with more negative values indicating favourable interactions, and is commonly estimated from MD trajectories using molecular mechanics energies with solvation and entropic corrections ^42,43^. In our study, negative ADS and ΔG_binding_ scores confirmed all ligands as promising Syt1 binders and thermodynamically favourable and spontaneous interactors, with ligand and protein RMSDs consistently below 4 Å throughout the MD trajectories (**Fig. 1D-F**), indicating stable binding. The SD patterns (**Table 1**) indicated that all complexes maintained structurally stable and thermodynamically favorable binding despite energetic heterogeneity and ligand-specific flexibility.

Correlation analysis of binding affinity scores with ligand physicochemical properties (**Fig. 1C**) revealed that Syt1–ligand bindings were strongly influenced by lipophilicity and hydrophobic interactions rather than polar contributions. This aligns with literature reporting that hydrogen bonds and hydrophobic contacts are low-energy, context-dependent contributors to thermodynamic stability, and that ligand-efficiency principles favour lipophilic interactions in hydrophobic pockets by reducing the desolvation cost of polar contacts ^38–40,44^.

The present study exhibited diverse subsets having certain overlaps (**Fig. 2A-H**) of interfacial residues including hotspots (**Fig. 2I**), with polar and nonpolar attributes, across ligand complexes. Bindings of all ligands were dominated by hydrophobic interactions. A limited subset of residues formed ligand-specific hydrogen bonds that might contribute to specificity and stabilization. Notably, K190–192, S235, and F349 formed hydrophobic contacts with six ligands (**Fig. 2A,B,E–H**), but not with MLT or EST (**Fig. 2C-D**). Literature shows that the residues K190-192, D232 and S235 in the C2A domain of Syt1 mediate calcium binding and membrane interactions, essential for neurotransmitter release upon calcium influx ^1^. Conversely, F349 in the C2B domain regulates oligomerization and clamp formation to control vesicle fusion timing ^45^. Especially, in the present study, D232 formed a hydrogen bond with the THI ligand (**Fig. 2G**). These residues did not appear as hotspots, yet their interactions with different ligands suggest their important roles in modulating Syt1 functions in Ca²⁺ sensing and regulating vesicular exocytosis.

### 4.2. Hotspot characterization: structural analyses

During interaction, protein interfaces assemble a conserved hydrophobic core of densely packed hotspots, defined by ASM-ΔΔG score ≥ 2 kcal.mol^-1^. These maximally solvent-excluded residues are surrounded by less energetic ‘null spots’ (ΔΔG < 2 kcal.mol^-1^). Together, these residues create a stabilizing ‘O-ring’ rim with low ‘solvent accessible surface area’ (SASA) as reported previously ^30,31^. In our study, 15 hotspots satisfied the ASM-ΔΔG threshold, with centroid distances to ligand Cα-atoms fluctuating between ≤4 Å and ≥8 Å during MD trajectories. Inter-residue contacts are reported to be within ≤4.5 Å for heavy atoms and ≤8 Å for Cα atoms ^46^, while α-carbon cutoffs ranging from 4 to 10 Å have been applied for residue connectivity and binding-pocket annotation ^47^. Therefore, hotspots in our study exhibited conformational adaptability, fluctuating within geometric thresholds, thereby maintaining stable Syt1–ligand binding.

PRDE-total values (**Fig. 2I**) revealed ligand-specific energetic patterns dominated by hydrophobic residues (methionine, leucine, phenylalanine, tryptophan). All fifteen hotspots showed negative mean PRDE-total values, indicating net stabilization, despite residue-dependent magnitudes and variability, with vdW interactions dominating over polar and non-polar solvation components (**Fig. 3C**). Literature indicates that a few binding residues disproportionately dominate free energy ^31^ and that protein interfaces exhibit context-dependent energetic sensitivity ^30^. Similarly, in this study, seven out of fifteen hotspots recurred across multiple ligands and most of them exhibited strong and ligand-dependent bindings dominated by vdW energy contributions. The remaining hotspots appeared to be mostly ligand-specific and showed predominant weak bindings due to counterbalance of vdW energetics by polar solvation energy (Eps). Collectively, the results indicated that binding energetics of hotspots were ligand- and context-dependent. Notably, hotspots clustered within the C2A, linker/N–C2B, and C2B regions, underscored their relevance in Syt-1 binding (**Fig. 2I**).

In water, dielectric screening weakens electrostatic interactions ^48^, while nonpolar molecular recognition are governed by vdW forces arising from the hydrophobic effects ^49^. These phenomena refer that solvation is governed by cavity-formation costs overriding solute-solvent interactions ^50^. Moreover, solvation free energy increases with solute size and surface complexity. As the irregular and concave nonpolar surfaces acquire higher cavitation costs, vdW-driven hydrophobic clustering diminishes solvent-exposed surface area coupled to enthalpy-entropy compensation ^49–52^. Conversely, polar groups stabilize solvent penetration and electrostatic hydration, thereby opposing hydrophobic association ^48,49^. Our results projected that vdW forces stabilized residues, while burial and solvent exclusion might drive enthalpy-entropy compensation, making polar hotspots behave hydrophobic-like dense packing at the expense of solvent accessibility rather than chemical identity.

### 4.3. Hotspot characterization: functional analyses

Syt1 dynamically alternates between open and closed conformations to regulate exocytosis. The close state, is characterized by a ∼108° rotation between C2A and C2B domains and stabilized by inter-domain salt-bridges between (a) D178 (Loop 1, C2A) and R388 (hydrophilic side of α-helix, C2B) and (b) R199/R233 (Loop 2/Loop 3, C2A) and D392 (hydrophilic side of α-helix, C2B). Additionally, interdomain hydrogen bonds between H237 and T406 restrain C2A-loop 3 from Ca²⁺ binding ^3,53^. The net negative charges of both C2-domains provide electrostatic repulsion to get rid of premature membrane fusion ^54^. In the open state, Ca^2+^ influx disrupts these interdomain interactions, enabling H237 and Y180 intra-domain bond within C2A that repositions loop 3 for Ca²⁺ binding ^3,53^. Ca²⁺ binding neutralizes negative charges of both C2 domains, providing electrostatic attraction that promotes membrane fusion, enables full SNARE complex zippering, and thereby drives exocytosis ^54^. Notably, the present study identified H237 and T406 as binding hotspots for the ligands MTF and THI respectively, both of which also serve as hotspots for MLT binding. Additionally, the residue Y180 appeared as one of the enlisted hotspots for THI binding. Therefore, binding of any of MLT, MTF and THI to Syt1 is predicted to modulate its inter- and intra-domain interactions, thereby regulating closed-to-open state conformation of Syt1. Both total (**Fig. 2I)** and components (**Fig. 3C**) of PRDE values also confirmed that ligand engagements of these hotspots were not only structurally plausible but also thermodynamically favourable, with distinct modes of adaptability across dominant, ligand-adaptive, and solvation-compensated hotspots.

It is reported that the stability of tripartite Syt1-(SNARE-complex)-complexin complex is mediated by Syt1-C2B interfacial contacts, including its (a) L394 with I203 of syntaxin-1A by hydrophobic coiled-coil-like interactions and (b) Q408 with D68 of complexin by hydrogen bonds ^55^. The present study identified common hotspots L394 for COR and EST and, Q408 for ARG, MTF and PLP. This result highlights the functional role of these ligands in modifying the tripartite interface interaction for vesicular anchoring and priming for exocytosis.

### 4.4. Desolvation effects of Syt1–ligand bindings

It is reported that non-polar solvation free energy depends nonlinearly on SASA, limiting surface-only descriptors ^56^. Alternatively, the V_SE_ measurement, decomposed into V_vdW_, internal void, and molecular surface volumes, provides a complementary geometric measure of protein packing and solvent inaccessibility relevant to stability and environmental responsiveness ^35^. Reductions in excluded volume upon association increase water translational entropy and favour binding, implying that increased excluded volume would oppose association ^57^. Tighter protein–ligand binding is associated with more negative binding volume changes, whereas weak binders show smaller or near-zero volume changes ^58^. Based on these thermodynamic principles, ligands with decrease in ΔV_SE_ (ARG, COR, EST, MTF, and THI) and increase in ΔV_SE_ (TIT, MLT, and PLP) represented enhancement and reduction of binding-site compaction, respectively (**Fig. 4**). Therefore, our results supported that ΔV_SE_ measurement was a structural depiction for hydration-mediated excluded-volume effects.

Notably, four hotspots with non-polar (F193, F377, M416) and neutral (Q408) side chains overlapped between PLP and ARG (**Fig. 2I**), yet exhibited divergent ΔV_SE_ profiles (**Fig. 4**), indicating that ΔV_SE_ discriminated ligands with shared structural hotspots but distinct thermodynamic effects. Divergent PRDE signatures across shared hotspots (**Fig. 2I**; **Fig. 3C**) underscored ΔV_SE_ as a discriminator of binding thermodynamics beyond geometry. The literature indicates that protein-ligand binding is governed by a non-additive interplay of thermodynamic determinants, which include (a) hotspot flexibility-mediated conformational entropy changes that modulates affinity and stability and (b) global solvation dynamics underlying enthalpy-entropy compensation ^59,60^. Notably, the ΔV_SE-overall_ of Syt1 had been increased across all ligand complexes. This reflected global volumetric redistribution and surface reshaping culminating in balanced local interfacial stabilization and solvation-driven conformational flexibility.

### 4.5. Linear versus non-linear correlations of MD features

Protein-ligand binding emerges from cooperative and nonlinear coupling of flexibility, entropy, and solvent reorganization ^60,61^, with desolvation and heterogeneous hydration dominating free energy landscapes ^62–64^. Quasi-linear trends appear only within restricted principles such as charge-driven solvation ^65^ or apolar burial ^62^. The present study revealed coexisting linear and nonlinear relationships among affinity (ΔG_binding_), structural flexibility (RMSDs), and solvation adaptability (ΔV_SE_) underlying binding stability (**Fig. 5**). Linear correlations across parameters captured thermodynamic-structural principles in which ΔG_binding_ covaried with conformational changes, whereas KPCA-IG uncovered a distinct nonlinear hierarchy of feature importance. Methodologically, KPCA-IG identifies dominant nonlinear features via gradient-assisted decomposition of kernel similarities and eigen decomposition of the kernel matrix ^37^. Accordingly, kPC1 dominance in feature space (**Fig. 5B**) reflected a hierarchical nonlinear organization in which collective solvent and conformational variability (ΔV_SE_, RMSD), rather than ΔG_binding_ alone, governed the principal axis of binding stability.

Linear positive correlation (**Fig. 5A**) showed that ligands achieving greater hydrophobic burial at the hotspot-enriched interface (i.e. negative ΔV_SE-hotspot_) enhanced binding affinity (negative ΔG_binding_). This agrees with reports that apolar surface burial strengthens protein-ligand affinity ^62^. Concomitantly, the strong negative correlation (**Fig. 5A**) revealed that tighter local packing (negative ΔV_SE-hotspot_) coupled with global protein expansion (ΔV_SE-overall_) indicated compensatory desolvation characteristic of induced-fit adaptable binding. Such adaptability reflects protein dynamics that balance global stability and functional flexibility within low-conflict energy landscapes ^66^, while highly disordered regions retain energetic conflicts to enable conformational diversity and partner specificity through suboptimal interactions ^67^.

Further, KPCA-IG scores (**Fig. 5B, right panel**) indicated that global desolvation (ΔV_SE-overall_) outweighed local desolvation (ΔV_SE-hotspot_), revealing nonlinear cooperative induced-fit effects that complemented linear energetics. Reports of ligand-induced local rigidification coupled to distal flexibility ^61^ and nonlinear solvent reorganization ^65^ support a globally adaptive binding mechanism. Consistently, heterogeneous hydration and protein conformational entropy contribute substantially to affinity differences beyond ΔG_binding_ alone ^60,63,64^. Thus, in our study, strong binding arose from linear affinity gains via hotspot-mediated hydrophobic burial coupled to cooperative induced-fit local rigidification, distal adaptation, and global desolvation. Further, RMSD analyses showed that binding-site optimization was achieved through selective conformational flexibility.

The trend in moderate linear correlation (**Fig. 5A**) associated (a) stronger binding affinity with minimal protein backbone rearrangement (ΔG_binding_ vs. RMSD_protein backbone_) and (b) ligand flexibility with docked geometry through local internal adjustments (RMSD_ligand_ vs. pose deviation), indicated that stable complex formation was achieved without global structural remodeling. This result aligns with evidence that the stable protein-ligand binding is accommodated by minimal backbone rearrangement with localized sidechain and solvent-mediated adjustments ^61^, while ligand pose stability is maintained primarily through internal conformational flexibility rather than rigid-body displacement ^68^.

KPCA-IG analysis (**Fig. 5B, right panel**) ranked structural contributions to binding stability as RMSD_ligand_> RMSD_complex(docked vs. post–MD)_>RMSD_protein-sidechain_>RMSD_protein-backbone_, identifying ligand conformational flexibility as the dominant driver of productive binding, followed by pose stabilization and limited protein restructuring. Literature indicates that affinity is governed primarily by ligand adaptability and accurate pose retention ^69^, while binding is refined by small, localized sidechain motions optimizing interactions and improving docking accuracy, consistent with reduced entropic costs ^70^, and sidechain heterogeneity provides compensatory flexibility elsewhere, preserving affinity and specificity ^61^. Taken together, our results indicate that binding stability was achieved through ligand-driven pose retention and localized sidechain optimization, rather than backbone rearrangement.

Collectively, our correlation results revealed a dual interpretation of binding stability. Linear correlations linked ΔG_binding_ to hotspot desolvation and structural stability for rapid affinity assessment. Nonlinear KPCA-IG revealed that ligand-driven solvation changes and conformational flexibility dominated productive binding. Strong affinity occurred when hydrophobic burial improved ΔG_binding_ and ligand flexibility preserved the docked pose through cooperative induced-fit adaptations, demonstrating that binding stability was governed by cooperative structural and hydration effects beyond ΔG_binding_ alone.

### 4.6. Ligand ranking based on correlations of MD features

The kPC1 and kPC2 plots indicated the balanced multidimensional variance in ligand space (**Fig. 5B, left panel**) with cooperative contributions of internal flexibility, solvent-coupled structural adjustment and pose retention. Accordingly, KPCA-IG rankings of ligands (**Fig. 5B, right panel**) integrated seven MD-features viz. ΔG_binding_ and RMSDs (**Table 1**), ΔV_SE_ values (**Fig. 4**) and pose deviation (**Fig. 3A**), rather than a single dominant determinant, to discriminate adaptive from rigid and burial-dominated binding states.

PLP and MLT exhibited solvent-efficient hotspot expansion and balanced structural variance that provided high nonlinear importance through adaptability-driven binding in complementary ways. PLP achieved moderate ΔG_binding_ through (a) strongly positive ΔV_SE-hotspot_ with minimal ΔV_SE-overall_, (b) restrained backbone yet adaptive side chains, and (c) controlled ligand fluctuations with progressive pose refinement, therefore showing localised adaptability with maximal nonlinear efficiency. In contrast, MLT attained comparable ΔG_binding_ with collective and stabilized interactions via ligand-protein-solvent network-based adaptability to sustain high nonlinear importance.

TIT binding generated a distinct MD profile viz. hydrophobic-driven interfacial expansion (positive but limited ΔV_SE-hotspot_) with moderate global solvent reorganization (ΔV_SE-overall_) and maximal backbone RMSD with subtle structural adjustments. Additionally, a hydrogen bond with V413 (Fig. 2H) might stabilize TIT pose while forming hydrophobic hotspot engagement without rigid locking.

Therefore, TIT achieved position alongside PLP and MLT in KPCA-IG ranking due to this anchoring modulated adaptability, despite its weaker ΔG_binding_ status.

MTF and COR exhibited hydrophobic interfacial burial (negative ΔV_SE-hotspot_) and diffuse solvent-mediated global compensation (elevated ΔV_SE-overall_) that projected passive burial-dominated states of their bindings. MTF showed pronounced ligand flexibility (highest RMSD_ligand_), restrained protein motion and a stable binding pose (lowest RMSD_complex(docked vs. post-MD)_) that yielded ligand-driven compensation rather than interfacial reorganization, consistent with its favourable ΔG_binding_ status. Conversely, COR binding represented a more rigid (low RMSD_ligand_) variant that induced moderate protein RMSDs and formed a stable complex with moderate pose deviation. However, the restricted dynamic-solvent (ΔV_SE_) responsiveness constrained nonlinear variance in both cases, placing them in the mid-tier of KPCA-IG importance.

ARG showed extreme hydrophobic-induced interfacial burial (highest negative ΔV_SE-hotspot_) with excessive global desolvation (largest ΔV_SE-overall_), constrained protein dynamics (backbone), substantial RMSD_ligand_ and stable pose convergence (low RMSD_complex(docked vs. postMD)_), therefore indicated adaptability-restricted coupling of ligand-protein-solvent during binding. Additionally, multiple hydrogen bonds (K182, N381, W404) might stabilize ARG binding by structural locking at the initial stage. Collectively, ARG achieved the lowest position in KPCA-IG ranking, depicting it as thermodynamically strong but mechanistically inefficient binder. Notably, both MTF and ARG exhibited comparable protein RMSDs (backbone and sidechain), while their KPCA-IG rankings differed due to their adaptive mechanisms, particularly ligand status (flexibility versus rigidity) within stable pose, which resulted in differential dynamic ligand-protein-solvent coupling.

Both THI and EST, like ARG, were dominated by polar-locked hydrogen bondings along with hydrophobic-induced interfacial burial (negative ΔV_SE-hotspot_). The specific hydrogen bonds viz. D232 for THI (**Fig. 2G**) and Q353 for EST (**Fig. 2C**) provided localized polar anchoring in both cases, alongside hydrophobic contacts. Additionally, variation in the magnitude of both hydrophobic burial and pronounced global solvent desolvation (high ΔV_SE-overall_) yielded minimal nonlinear KPCA-IG contribution for THI and EST.

THI exhibited rigidity (very low RMSD_ligand_) with adjusted protein motions (low RMSD_sidechain_ and high RMSD_backbone_) and formed pose-locked complex (moderate RMSD_complex(docked vs. post–MD)_). Together, THI showed limited dynamic ligand-protein-solvent coupling. Despite having modest ΔG_binding_, THI reflected non-adaptive burial with substantial solvent cost, which could place THI in a position immediately below ARG in KPCA-IG ranking. Additionally, THI and COR displayed comparable RMSDs and similar pose stability. COR retained greater dynamic responsiveness to interfacial and solvent reorganization, yielding a higher KPCA-IG rank than THI.

EST showed negligible hotspot engagement (near-zero ΔV_SE-hotspot_) and energetically costly diffuse global solvent displacement (extremely high ΔV_SE-overall_) that provided solvent-inefficient regime. Rigid EST binding (lowest RMSD_ligand_) and its lagging non-cooperative pose adjustment (highest RMSD_complex(docked vs. post-MD)_) were accompanied by constrained protein dynamics. These features yielded the weakest ΔG_binding_, with rigid structural stabilization and cooperative ligand-protein-solvent adaptability, placing EST in lowest KPCA-IG rank.

These results highlighted that stable binding required a balance of rigidity, ligand mobility, solvent-efficient induced-fit adaptation, and enthalpy–entropy compensation, rather than affinity alone. Literature reports indicate that effective ligands combine pose retention, structural compatibility and dynamics ^71,72^, a balance of controlled desolvation and hydration-aware cooperativity ^63,71,72^, and thermodynamic compensation ^63,73^.

In conclusion, the novelty of this study resides in a thermodynamic–structural–solvation framework showing that productive Syt1–ligand binding is governed by desolvation, molecular flexibility and induced-fit adaptability rather than ΔG_binding_ alone, enabling predictive classification into adaptive, passive, and locked modes. Integrating ASM-ΔΔG with PRDE scores establishes a unified framework in which hotspot stabilization is consistently supported by negative per-residue energies, thereby reinforcing this classification. These predictions resolve C2A- or C2B-biased hotspot engagement that couples ligands to Ca²⁺-dependent conformational switching or SNARE–complexin regulation, and further identify auxiliary non-hotspot interactions that tune Ca²⁺ sensing, membrane association, and fusion-clamp control. Together with nonlinear importance ranking, this work identifies Syt1 as a multiligand neuromodulatory scaffold and establishes a dual-criteria strategy, combining functional hotspot engagement and nonlinear adaptability, for presynaptic modulation and neurological drug discovery.

## Acknowledgements

The present study was funded by the Department of Biotechnology (DBT), Ministry of Science and Technology, Government of India, under BINC (Bioinformatics National Certification) scheme [No.: BT/BI/10/078/2014]. We are thankful to Professor Pritha Mukhopadhyay, coordinator of CPEPA-UGC centre [“Centre for Electrophysiological and Neuro-imaging studies including Mathematical Modelling” (CPEPA) through “University Grants Commission” (UGC)], under the University of Calcutta, India. We gratefully acknowledge Professor David A. Case (Department of Chemistry & Chemical Biology, Rutgers University) and Alma Rosa Agorilla (AMBER Software Administrator, University of California, San Francisco) for approving the academic fee waiver and providing access to the AMBER18 molecular dynamics software. The authors sincerely thank Dr. Roman Laskowski (European Bioinformatics Institute, European Molecular Biology Laboratory, Hinxton, UK) for his support and granting access to the LigPlot+ software.

## Funding

The Senior Research Fellowship grant was funded by the Department of Biotechnology (DBT), BINC (Bioinformatics National Certification), Ministry of Science and Technology, Government of India, to Suvojit Hazra (Grant No.: BT/BI/10/078/2014).

## CRediT authorship contribution statement

**Suvojit Hazra:** Conceptualization, Data Curation, Formal analysis, Investigation, Methodology, Resources, Software, Validation, Visualization, Writing - Original Draft. **Basant K. Tiwary:** Validation, Investigation, Formal analysis, Methodology, Resources, Writing - Review & Editing, Project administration. **Alok Ghosh Chaudhuri:** Conceptualization, Validation, Writing - Review & Editing. **Nilkanta Chakrabarti:** Conceptualization, Investigation, Formal analysis, Methodology, Resources, Validation, Visualization, Writing - Review & Editing, Supervision, Project administration.

## Data availability

All data associated with this study are present in the paper.

## Declaration of competing interests

The authors declare that they have no known competing financial interests or personal relationships that could have appeared to influence the work reported in this paper.

## Notes

### Competing Interest Statement

The authors have declared no competing interest.

